# Fine-mapping of Parkinson’s disease susceptibility loci identifies putative causal variants

**DOI:** 10.1101/2020.10.22.340158

**Authors:** Brian M. Schilder, Towfique Raj

**Affiliations:** Nash Family Department of Neuroscience & Friedman Brain Institute, Icahn School of Medicine at Mount Sinai, New York, NY, United States of America; Ronald M. Loeb Center for Alzheimer’s disease, Icahn School of Medicine at Mount Sinai, New York, NY, United States of America; Department of Genetics and Genomic Sciences, Icahn School of Medicine at Mount Sinai, New York, NY, United States of America; Icahn Institute for Data Science and Genomic Technology, Icahn School of Medicine at Mount Sinai, New York, NY, United States of America; Estelle and Daniel Maggin Department of Neurology, Icahn School of Medicine at Mount Sinai, New York, NY, United States of America

**Author notes:** Corresponding Author Towfique Raj.

**Keywords:** Genome-wide association, Expression quantitative trait loci (eQTL), Neurodegeneration, Neurogenetics, Bioinformatics, Fine-mapping

## Abstract

Recent genome-wide association studies have identified 78 loci associated with Parkinson’s Disease susceptibility but the underlying mechanisms remain largely unclear. To identify variants likely causal for disease risk, we fine-mapped these Parkinson’s-associated loci using four different statistical and functional fine-mapping methods. We then integrated multi-assay cell-type-specific epigenomic profiles to pinpoint the likely mechanism of action of each variant, allowing us to identify Consensus SNPs that disrupt *LRRK2* and *FCGR2A* regulatory elements in microglia, *MBNL2* enhancers in oligodendrocytes, and *DYRK1A* enhancers in neurons. Finally, we confirmed the functional relevance of fine-mapped SNPs using a suite of *in silico* validation approaches. Together, these results provide a robust list of likely causal variants underlying Parkinson’s Disease risk for further mechanistic studies.

## Introduction

Parkinson’s Disease (PD) is the second most prevalent neurodegenerative disease, occurring in 2-3% of individuals 65 years of age or older^1^. Many efforts have been made to better understand the biological mechanisms underpinning this disease in hopes of developing more effective treatments. Genome-wide association studies (GWAS) have offered insight into the molecular etiology of this debilitating disease^2–4^.

The largest PD GWAS to date recently identified 90 independent PD-associated variants distributed across 78 loci^4^, more than doubling the number of previously known PD-risk signals^3^. However, for any given locus the lead or tag single nucleotide polymorphism (SNP) may merely be correlated with the causal SNP(s) due to linkage disequilibrium (LD), thus limiting our ability to interpret the functional consequences of genetic variation through which they affect complex disease risk (e.g. PD)^5–7^. Fine-mapping is a category of methodological approaches that aim to prioritize putative causal variants^8–10^. There are currently a wide variety of tools that can perform statistical, functional, and trans-ethnic fine-mapping, each with their own sets of strengths and weaknesses. In order to robustly and efficiently fine-map PD loci, we developed *echolocatoR^11^*, an open-access R package that enables automated end-to-end fine-mapping across many loci, using multiple fine-mapping tools in a single R function (**Fig. 1**). We fine-mapped the majority of the PD loci using four different statistical and functional fine-mapping tools, which reduced the average number of candidate SNPs per locus from 4,981 to just 3. The resulting fine-mapped SNPs, especially the multi-tool Consensus SNPs, were greatly enriched for functional annotations compared to the lead GWAS SNPs. Furthermore, we identified many Consensus SNPs that are within cell-type-specific regulatory regions (e.g. active microglia enhancers), providing comprehensive and novel insights into PD molecular etiology.

**Fig. 1.**
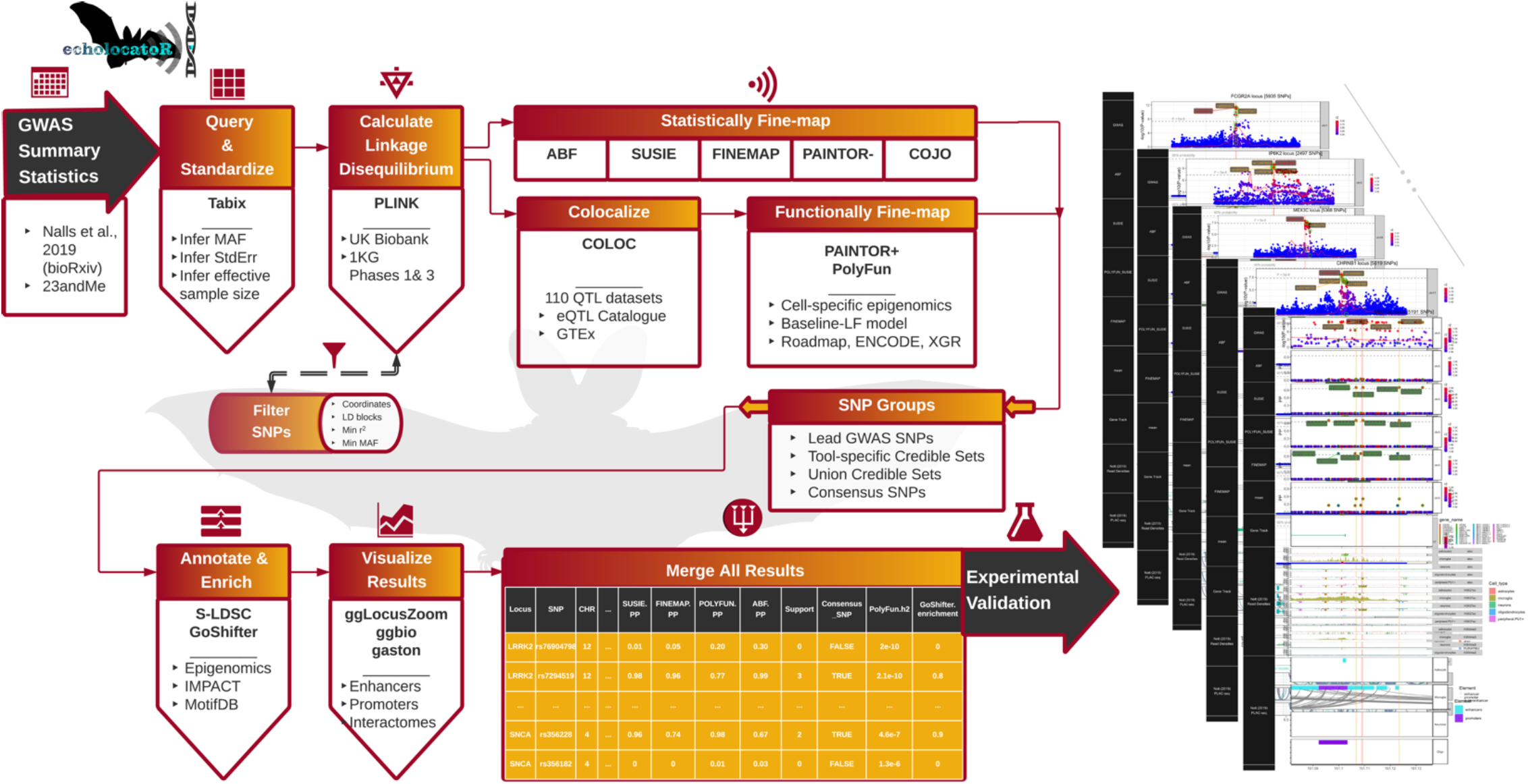
Outline of the study workflow. *echolocatoR* enables end-to-end fine-mapping using the following workflow: 1) Nalls et al.^4^ GWAS summary statistics imported, 2) locus-specific subsets are queried and standardized, 3) LD matrices are extracted from a European-ancestry subset of UK Biobank (UKB)^17,62,63^, or the 1000 Genomes Project (1KG), 4) statistical and functional fine-mapping are conducted across multiple tools, 5) SNPs are categorized, 6) enrichment for tissue- and cell-type-specific epigenomics, S-LDSC heritability, predictions of functional impact from several machine learning models (IMPACT, Basenji, DeepSEA), and motif disruption (*motifbreakR*), 7) all results are merged into a single table with one row per SNP, 8) high-resolution multi-track plots are generated for each locus.

## Results

### Fine-mapping of PD GWAS loci

After removing loci that fall within the HLA or Tau/17q21.31 regions (which were excluded due to the known complexity of their LD architectures)^12^, we present here fine-mapping results for 74/78 loci using four complementary fine-mapping tools: ABF^13^, FINEMAP^14,15^, SuSiE^16^, and PolyFun+SuSiE^16,17^. Several key SNP groups were compared here: 1) *GWAS lead*: the SNP with the smallest p-value in each PD GWAS locus (Nalls et al. 2019) *after* filtering steps (i.e. removing SNPs with MAF < 0.05 or are absent in the LD reference panel, and considering all SNPs within ± 1Mb of the original pre-filtered GWAS lead SNP), 2) *[fine-mapping tool name] CS*_*95%*_: tool-specific 95% credible sets, generally defined as having ≥ 95% probability of being causal for the phenotype (i.e. PD), 3) *UCS*: Union Credible Set SNPs defined as the union of all tool-specific CS_95%_;, and 4) *Consensus*: SNPs in the CS_95%_; of at least two fine-mapping tools.

Using our multi-tool fine-mapping strategy, we identified UCS and Consensus SNPs in 100% of these 74 loci. In total, there were 598 UCS SNPs (8.1 per locus on average) and 190 Consensus SNPs (2.6 per locus on average). The proportion of loci that each tool was able to identify at least one CS_95%_ SNP varied considerably (**Fig. 2**). For example, ABF (a relatively simple fine-mapping model that can only assume one causal SNP/locus) was only able to identify a CS_95%_ in eight loci, whereas SuSiE (which can assume multiple causal SNPs) produced CS_95%_ in all loci. We also note that the SNP-wise posterior probabilities (PP) of PolyFun+SuSiE and SuSiE are expectedly much more highly correlated with one another than those of ABF or FINEMAP (**Fig. S1g**).

**Fig. 2.**
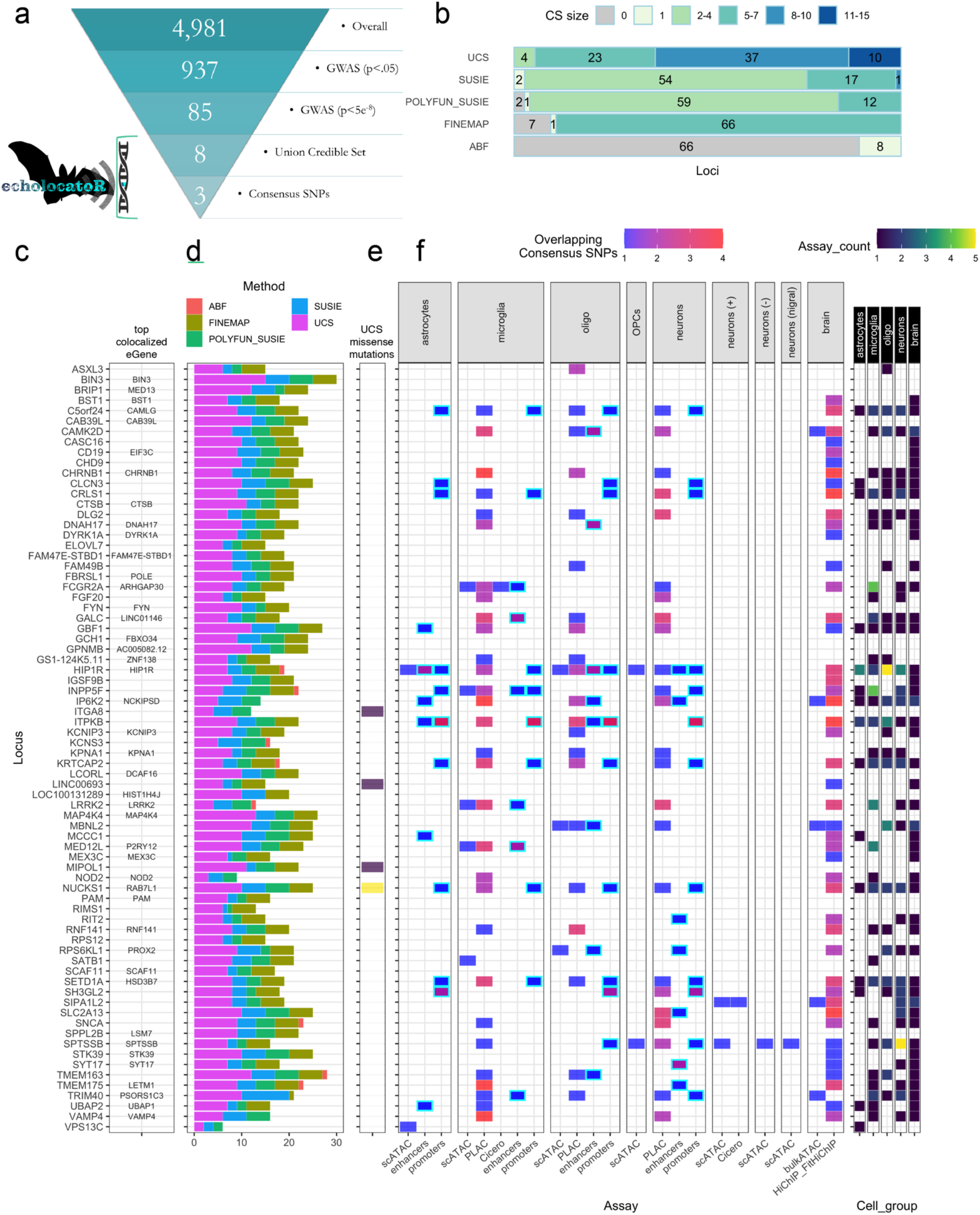
Summary of all fine-mapped loci. **a**, The average number of potentially causal SNPs per PD-associated GWAS locus was reduced from 4,981 to just 3 Consensus SNPs. **b**, The number of loci in which each fine-mapping tool identified a 95% Credible Set (CS_95%_) of a given size. These counts are also shown for the Union Credible Set (UCS) of all CS_95%_ together. **c**, Rows are names of each locus as designated in the Nalls et al.^4^ GWAS, while “top colocalized eGene” displays the gene that showed the highest colocalization probability across all tested eGenes in all eQTL Catalogue datasets (after filtering spurious RP11-genes). **d**, The number of SNPs in each tool-specific CS_95%_, as well as UCS size. **e**, The number of UCS SNPs that were missense mutations. **f**, The number of Consensus SNPs that fell within different cell type-specific epigenomic annotations. Nott et al.^18^ data includes enhancers or promoters called from peaks across multiple assays in the as well as overlap with PLAC-seq co-accessibility anchors (PLAC). Data from Corces et. al (2020) includes single-cell ATAC-seq peaks (scATAC), co-accessibility anchors called from the scATAC-seq data using the tool Cicero (Cicero), as well as peaks called from ATAC-seq in bulk brain tissue (bulkATAC) and co-accessibility anchors from HiChIP-FitHiChIP in bulk brain. (OPCs = oligodendrocyte progenitor cells, neurons (+) = excitatory neurons, neurons (-) = inhibitory neurons, neurons (nigra) = dopaminergic neurons from the substantia nigra. **g**, The number of assays in which at least one Consensus SNP overlapped with the annotation, aggregated by cell type (or bulk brain). This highlights the most relevant cell type(s) in each locus.

When checking for overlap with cell-type-specific epigenomic annotations^18,19^, we found that 66/74 loci (89.2%) had at least one overlapping UCS SNP and 54/74 loci (73%) loci had at least one overlapping Consensus SNP (**Fig. 2f**). We also checked for exclusivity amongst brain cell-type-specific epigenomic signatures (astrocytes, microglia, neurons and oligodendrocytes in any assay), and found that Consensus SNPs overlapped with annotations in only one cell-type in 14 loci, two cell-types in 12 loci, three cell-types in 10 loci, and four cell-types in 9 loci. UCS SNPs overlapped with enhancers in 42 loci and promoters in 20 loci, while Consensus SNPs overlapped with enhancers in 21 loci and promoters in 11 loci. See the *In silico validation* section below for quantitative enrichment analysis results.

Mismatch between the population compositions of the LD reference panel and the original GWAS, as well as insufficient LD reference panel size, can have drastic effects on fine-mapping results^20^. We therefore repeated the fine-mapping pipeline on all PD loci using the European ancestry subset of 1000 Genomes Phase 3 (1KG; n = 503 samples) as the LD reference panel instead of UKB (n = 337,000 samples). Due to differences between 1KG and UKB, not all SNPs were present in both panels. Thus, only 62/74 (83.8%) of loci had the same lead SNP between panels. We next compared fine-mapping UCS using LD from each panel, and found that all loci had at least one shared UCS SNP between panels. The multi-tool mean PP was highly correlated between LD panels (Spearman rho = 0.65, p < 2.2×10^-16^), though see **Fig. S1a-g** for plots of inter-tool variation. In 46/74 (62.2%) of loci, at least one Consensus SNP overlapped between LD panel results. Despite this, when we specifically checked the loci that we show in multi-track plots (LRRK2, **Fig. 3a**; MBNL2, **Fig. 3b**; DYRK1A, **Fig. 4a**; FCGR2A, **Fig. 4b**) we found that all Consensus SNPs using the UKB panel were also Consensus SNPs using the 1KG panel, affirming that the Consensus SNPs in these example loci are robust and reproducible.

**Fig. 3.**
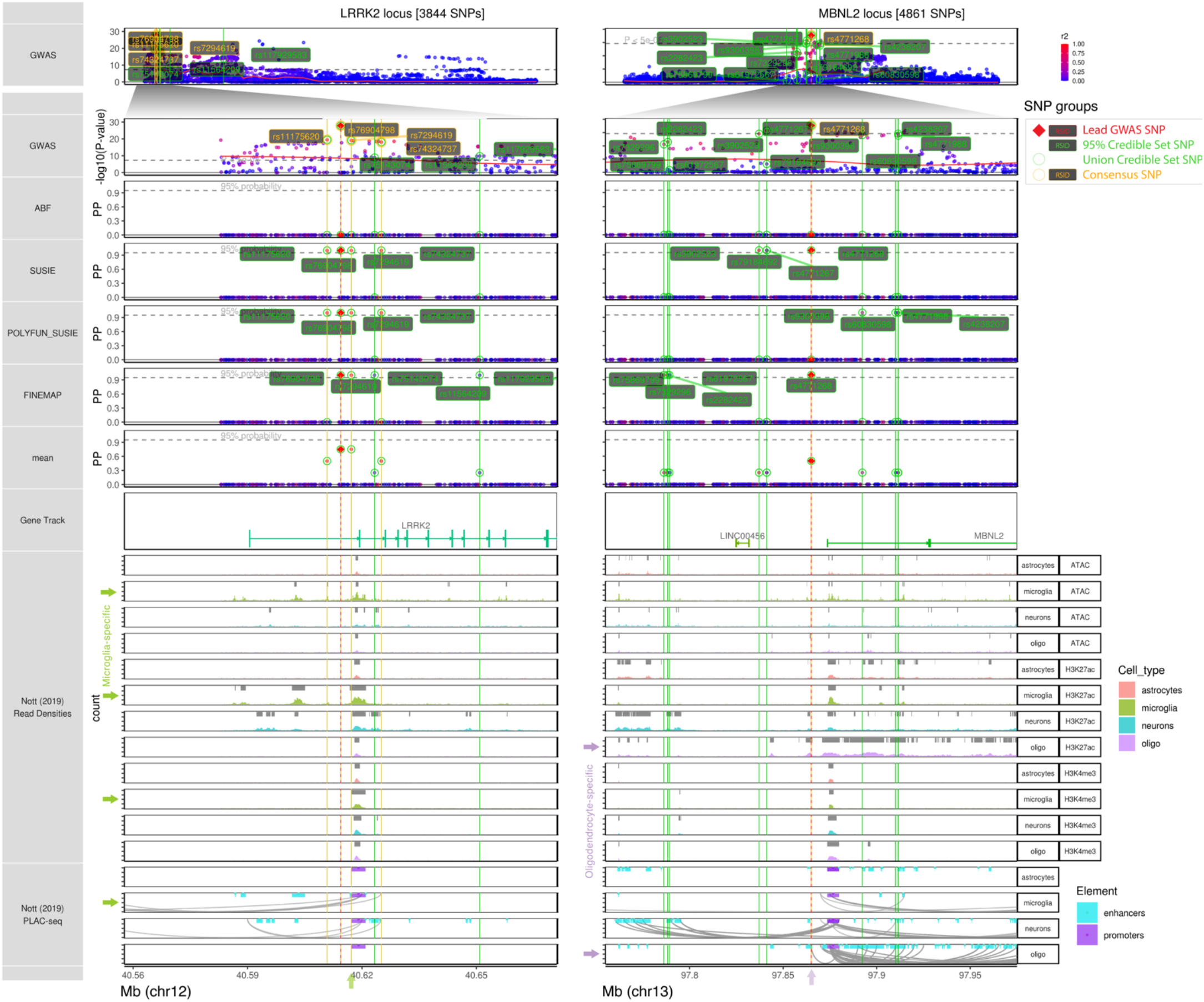
Fine-mapping of LRRK2 and MBNL2 loci. Multi-track plots within two PD-associated loci. **a**, LRRK2 contains Consensus SNPs that are common in the population and overlap with microglia-specific epigenomic peaks (indicated by green arrows). Specifically rs7294619 (which is not the lead SNP) falls within a microglia-specific enhancer, implicating a cell-type-specific mechanism of PD risk. **b**, MBNL2 contains a single Consensus SNP (rs4771268), which happens to also be the lead GWAS SNP, that overlaps with oligodendrocyte-specific epigenomic peaks (indicated by purple arrows). It also falls within an enhancer, with direct interactions (as indicated by PLAC-seq arches) with the MBNL2 promoter. The following tracks are shown (track labels in grey boxes on the left, subtrack labels in white boxes on the right): **GWAS**: -log10(p-value) from the Nalls et al.^4^ PD GWAS zoomed out to show the entire locus. Below is the same GWAS data but zoomed into 10x to better show the fine-mapped SNPs. (red diamond = lead GWAS SNP, green label = UCS SNPs, gold label = Consensus SNPs). **ABF, SUSIE, POLYFUN_SUSIE, FINEMAP**: Fine-mapping results from four different tools, with the posterior probability (PP) that each SNP is causal as the y-axis. (red diamond = lead GWAS SNP, green label = tool-specific CS_95%_ SNPs, green circles = UCS SNPs). **Mean:** Per-SNP PP averaged across all fine-mapped tools. **Gene Track:** Gene model of one of the LRRK2 transcripts (other transcripts not shown for simplicity). **Nott (2019) Read Densities:** Histograms of cell-type-specific assays (oligo = oligodendrocytes). Called peaks are indicated by grey bars. **Nott (2019) PLAC-seq**:PLAC-seq interactome data as well as enhancers and promoters called by the original authors.

**Fig. 4.**
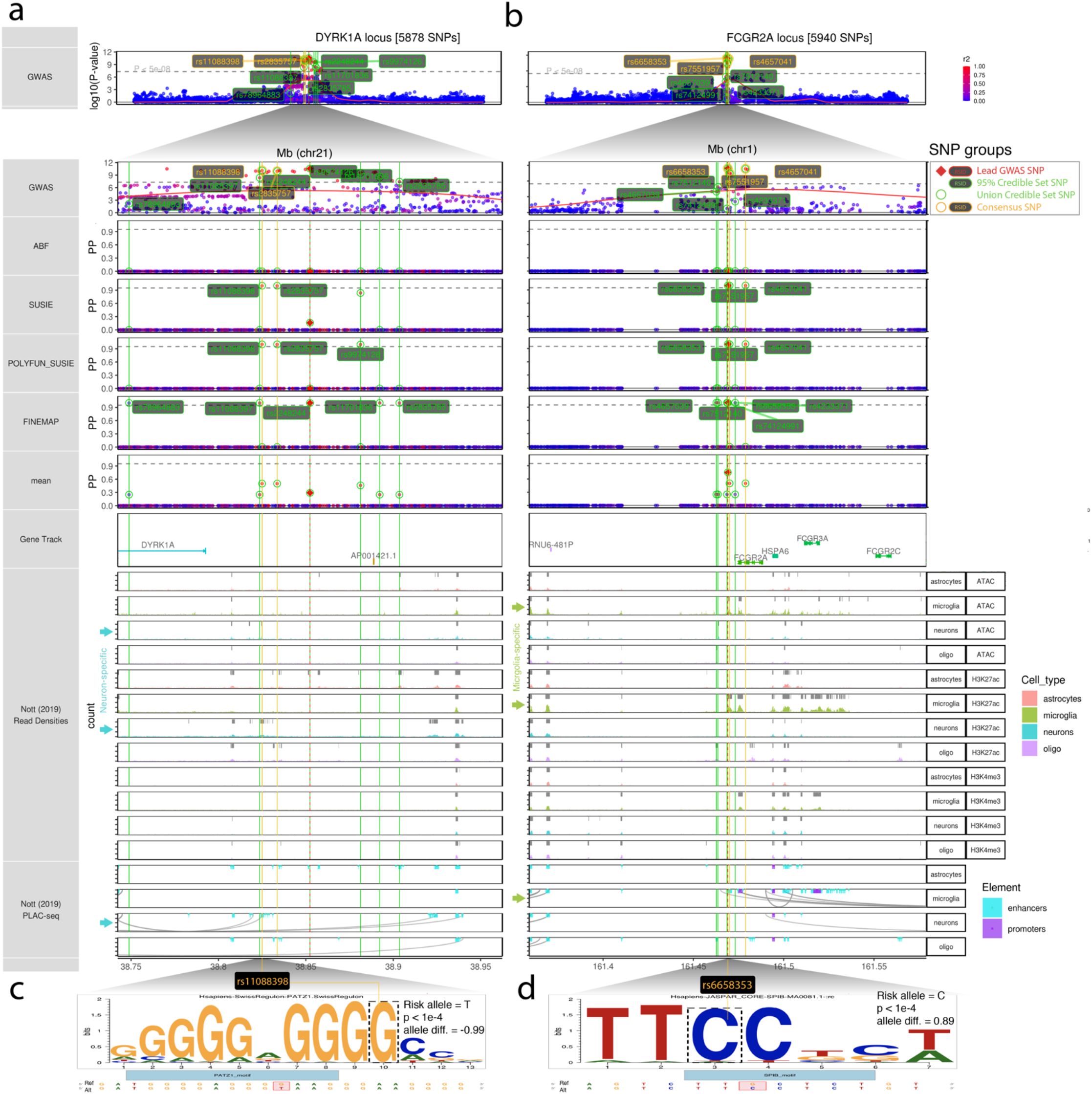
Fine-mapping of DYRK1A and FCGR2A loci. Multi-track plots within two PD-associated loci. **a**, The DYRK1A locus contains common Consensus SNPs that exclusively overlap with neuron-specific epigenomic peaks (indicated by blue arrows). Specifically rs11088398 (which is not the lead SNP) falls within a neuron-specific enhancer which has direct interactions with an upstream DYRK1A promoter. c, *motifbreakR* results indicating that rs11088398 strongly disrupts a PATZ1 transcription factor binding motif (TFBM) which remains highly significant even after accounting for background effects and multiple testing correction (p < 1×10^-4^). **B**, The FCGR2A locus contains three Consensus SNPs, all of which exclusively overlap with microglia-specific epigenomic peaks (indicated by green arrows). Specifically, rs665835 falls within a microglia-specific enhancer that is predicted to strongly disrupt binding in a *SPIB* TFBM (p < 1×10^-4^). It also falls within an enhancer, with direct interactions (as indicated by PLAC-seq arches) with the MBNL2 promoter. Within subplots **a** and **b**, the following tracks are shown (track labels in grey boxes on the left, subtrack labels in white boxes on the right): **GWAS** -log_10_(p-value) from the Nalls et al.^4^ PD GWAS zoomed out to show the entire locus. Below is the same GWAS data but zoomed into 10x to better show the fine-mapped SNPs. (red diamond = lead GWAS SNP, green label = UCS SNPs, gold label = Consensus SNPs). **ABF, SUSIE, POLYFUN_SUSIE, FINEMAP** Fine-mapping results from four different tools, with the posterior probability (PP) that each SNP is causal as the y-axis. (red diamond = lead GWAS SNP, green label = tool-specific CS_95%_ SNPs, green circles = UCS SNPs). **Mean** Per-SNP PP averaged across all fine-mapped tools. **Gene Track** Gene model of one of the LRRK2 transcripts (other transcripts not shown for simplicity). **Nott (2019) Read Densities** Histograms of cell-type-specific assays (oligo = oligodendrocytes). Called peaks are indicated by grey bars. **Nott (2019) PLAC-seq** PLAC-seq interactome data as well as enhancers and promoters called by the original authors.

To assess reproducibility, we also compared our FINEMAP results with those from a recent study that performed statistical fine-mapping (using FINEMAP) on the same PD GWAS meta-analysis dataset but using a TOPMed (n = 16,257 samples) as the LD reference panel^21^ (**Fig. S1h-m**). When comparing our UCS SNPs to a subset of the FINEMAP results provided by Grenn and colleagues^21^ (42 CS_95%_ SNPs across 18 loci), we found 10 overlapping SNPs, 6 of which were Consensus SNPs and 5 of which were also in our FINEMAP CS95%, across 9 loci (HIP1R, KRTCAP2, SH3GL2, SLC2A13, FAM47E-STBD1, TMEM175, TMEM163, CRHR1, and KCNS3). This limited concordance (10/42 SNPs = 23.8%) likely stems from several key methodological differences, including different LD panels and the fact that we specify a maximum of five causal SNPs for all loci whereas Grenn et al. first estimated the number of independent causal signals in each locus using the stepwise model selection procedure in GCTA-COJO^22^. Despite this, we observed moderate correlation between the SNP-wise FINEMAP posterior inclusion probabilities (PIP) from Grenn et al. and those of our FINEMAP analyses (Spearman rho = 0.68, p-value < 2.2×10^-16^, n = 6,391 overlapping SNPs; **Fig. S1k**), as well as with mean.PP in UCS SNPs only (Spearman rho = 0.68, p-value < 2.2×10^-16^, n = 10). This suggests that despite substantial methodological differences, there is moderate concordance in the pattern of results between studies.

### Examples of fine-mapped loci: LRRK2, MBNL2, DYRK1A and FCGR2A

To exemplify the utility of our fine-mapping approach, here we highlight results from four loci and further provide a web app for results in all loci: https://rajlab.shinyapps.io/Fine_Mapping_Shiny

Rare mutations within protein-coding domains of the *LRRK2* gene are frequently found in familial PD^23,24^. However, less is known about potential common causal variants at this locus (MAF > 1%). Here, we identified four common Consensus SNPs (MAF > 10%) within the LRRK2 locus, of which only rs7294619 (not the lead PD GWAS SNP) is within a cell-type-specific (microglia) enhancer (**Fig. 3a**). This confirms early independent reports of rs7294619 as a PD risk factor in smaller subpopulations^25^. eQTL colocalization tests further implicate *LRRK2* as the most functionally relevant gene in this locus in both microglia (coloc PP.H4 = 0.65, n = 93 individuals)^26^ as well as peripheral monocytes, a closely related myeloid cell-type, in multiple more well-powered datasets (BLUEPRINT: PP.H4 = 0.999, n = 554; Fairfax_2014: PP.H4 = 0.968, n = 1,372; Quach_2016: PP.H4 = 0.991, n = 969; **Fig. S2**)^27–29^. Specifically, rs7294619 is a significant eVariant that increases only *LRRK2* expression in microglia (p = 1.27×10^-6^, beta = 0.263)^26^. The involvement of *LRRK2* in microglia dysregulation aligns with substantial previous research implicating this gene in systemic and central nervous system (CNS) inflammation aspects of PD^23,24,30^ and other inflammatory diseases^31–33^.

Within the MBNL2 locus, we identified 11 UCS SNPs and just one Consensus SNP (rs4771268), which was also the lead GWAS SNP. rs4771268 overlaps with regulatory elements (specifically enhancers) exclusively active in oligodendrocytes, lending further support to recent studies that have indicated a more important role of oligodendrocytes in PD than previously suspected^18,34^. This oligodendrocyte-specific enhancer interacts with a downstream promoter of the gene *MBNL2* (see Proximity Ligation-Assisted ChIP (PLAC) -seq interactome tracks in **Fig. 3b**), supporting the hypothesis that *MBNL2* is likely to be the causal gene within this locus (though the eQTL colocalization tests could not identify any eGene for this region). We also conducted transcription factor binding motif (TFBM) analyses or the LRRK2 and MBNL2 loci using *motifbreakR* (**Fig. S3a-c**), although the proposed TFBM were less consistent across motif databases and we therefore caution readers to interpret these with caution.

At the DYRK1A locus, we identified two Consensus SNPs (rs11088398 and rs2835757, neither of which were the lead GWAS SNP) but only rs11088398 falls within a neuron-specific enhancer and has direct interactions with the upstream promoter of *DYRK1A*, as shown by multiple interactome assays (**Fig. 4a)**. Previously, Grenn et al.^21^ used FINEMAP to nominate rs2248244 (the lead SNP) and rs11701722 as putative causal variants. Despite the aforementioned methodological differences, rs2248244 also appeared in our UCS. However, in contrast to rs11088398, rs2248244 does not fall within any cell-type-specific epigenomic peak or regulatory annotation in the datasets explored here. While multiple variants within this locus may influence PD risk via different mechanisms, we find the functionally mechanistic explanation for rs11088398 particularly compelling. In further support of this, analysis of this region using *motifbreakR^35^*, which searches a comprehensive database of positional weight matrices (PWM) and applies significance-based background correction, revealed that rs11088398 is within a *PATZ1* transcription factor binding motif (TFBM), as well as *KLF5* and *MAZ* TFBM to a lesser extent (p < 4.15×10^-4^, allele diff. > -0.785), and that the PD risk allele (G→T) is predicted to strongly decrease binding affinity relative to the reference allele (p < 1×10^-10^, allele diff. = -0.980; **Fig. 4b**). This association remained highly significant even after background correction and stringent Bonferroni multiple-testing correction (q < 1×10^-10^). *PATZ1* plays an important role in embryonic development and neurogenesis (e.g. in midbrain), is expressed in neurons (and not glia) (**Fig. S3d)** and its downregulation is associated with premature senescence in mouse models, cell cultures, and human brain tissue^36,37^. Lastly, *DYRK1A* is the top eQTL-nominated gene from our colocalization analysis, which aligns with prior eQTL-based gene nominations in this locus^21^. It is expressed in all brain cell types, but more so neurons than glia (**Fig. S3e**).

Lastly, the gene Fc Fragment Of IgG Receptor IIa (*FCGR2A*) codes for an IgGFc receptor protein expressed on the surface of immune response cells and is important for phagocytosis and debris clearing^38^. Our fine-mapping analyses revealed three Consensus SNPs, two of which (rs6658353, rs7551957) fell within a microglia-specific enhancer (to the exclusion of other cell-type-specific epigenomic peaks) in the FCGR2A locus. While the Consensus SNP did fall within multiple interactome assay anchors (PLAC-seq, Cicero, HiChIP-FitHiChIP)^18,19^, the data did not associate it with a specific gene. *FCGR2A* was nevertheless the closest to the enhancer that the Consensus SNPs were located in. Furthermore, the Consensus SNP rs6658353 falls within a *SPI1* (a well-established transcriptional regulator in microglia) TFBM, and its PD risk allele (G→C) greatly disrupts its binding (**Fig. 4d**; p < 1×10^-4^, allele diff. = 0.89). *FCGR2A* itself is very highly expressed in both microglia and macrophages (**Fig. S3f**), as is *SPI1* (**Fig. S3g**).

### *In silico* validation

To efficiently verify the functionality of the fine-mapped SNPs, we employed an *in silico* validation strategy across a diverse set of functional annotations (i.e. validation datasets), including: 1) heritability enrichment scores derived from the L2-regularized S-LDSC regression^39–41^ step in PolyFun^17^, 2) probability scores from IMPACT, an elastic net logistic regression model that integrates 503 cell-type-specific epigenomic annotations to predict each variant’s functional impact in the context of a particular tissue and cell type, primarily through the disruption of transcription factor (TF) binding motifs^42^, 3) per-variant p-values from tests of genotypes impact on gene expression from the survey of regulatory elements (SuRE) massively parallel reporter assay (MPRA)^43^, as well as predictions from deep learning models, Basenji^44^ and DeepSEA^45^, trained on blood-, brain- or non-tissue-specific annotations^46^.

Specifically, we tested two hypotheses within each validation dataset (H1 and H2). H1: fine-mapped (e.g. *UCS, Consensus*) SNPs have greater functional impact, and thus are more likely to impact disease risk, than *GWAS lead* SNPs. H1 was tested using pairwise Wilcoxon rank-sum tests on per-Locus means of each SNP group. H2: fine-mapped SNPs more frequently have greater functional impact than randomly selected SNPs than do GWAS lead SNPs. H2 was tested using 10,000 boot-strapped iterations to compare functional annotations from each resampled SNP group to those of randomly sampled SNPs. In addition to the four main SNP groups defined above, we also compared SNP groups called without any functional fine-mapping (i.e. PolyFun+SuSIE) to avoid circularity, as some of the validation datasets (i.e. IMPACT, Basenji, DeepSEA) used annotations that were also used in the PolyFun baseline model. These additional SNP groups were: 5) *UCS (-PolyFun)*, and 6) *Consensus (-PolyFun)*. All p-values listed below are post-adjustment, unless otherwise specified. While an in depth analysis of inter-tool performance is beyond the scope of this study, we do provide extended *in silico* validation results comparing tool-specific CS95%(**Figs. S4 & S5**). See **Methods** and **Supplementary Methods** for extended details on these analyses.

Relative to GWAS lead SNPs, UCS SNPs had significantly greater h^2^ enrichment scores (H1 p = 3.2×10^-6^, H2 p = 6.2×10^-88^, mean = 1.46; **Figs. 5a**), IMPACT probability scores (H1 p = 0.0053, H2 p = 3.7×10^-238^, mean = 0.69; **Fig. 5b**), and functional impact probability in all 28 deep learning model-tissue-assay combinations (H1 p < 0.05; H2 p < 2.7×10^-303^, mean = 0.0056; **Fig. 5d**). The significant gain in predicted functionality remained true for UCS (-PolyFun) SNPs: h^2^ enrichment (H1 p = 1×10^-5^, H2 p = 7.3×10^-132^, mean = 1.37), IMPACT probability scores (H1 p = 0.046, H2 p = 2.2×10^-308^, mean = 0.68), functional impact probability in all 28 deep learning model-tissue-assay combinations (H1 adj. p < 0.05; H2 norm. p = 2.7×10^-303^, mean = 0.0056; **Fig. 5d**). Likewise, Consensus SNPs also showed significantly greater functional impact than GWAS lead SNPs: h^2^ enrichment (H1 p = 0.00052, H2 p = 9.3×10^-239^, mean = 1.68), IMPACT probability scores (H1 p = 0.00031, H2 p = 2.2×10^-308^, mean = 0.74), functional impact probability in 18/28 deep learning model-tissue-assay combinations (H1 p < 0.05; H2 p = 2.7×10^-303^, mean = 0.0053; **Fig. 5d**). While the H1 test of SuRE MPRA did not show significant differences between SNP groups, the bootstrapping test revealed that there was indeed significantly greater impact on gene expression in UCS, UCS (-PolyFun), and Consensus SNPs (H2 p = 2.7×10^-303^). This is perhaps due to the ability of the bootstrapping approach to approximate the true underlying distribution, and is more robust to sampling error. While Consensus SNPs were not significantly greater than UCS or UCS (-PolyFun) in any of the H1 or H2 tests, they still have fewer SNPs on average (~3 per locus) making them better suited for follow up wet lab experiments.

**Fig 5.**
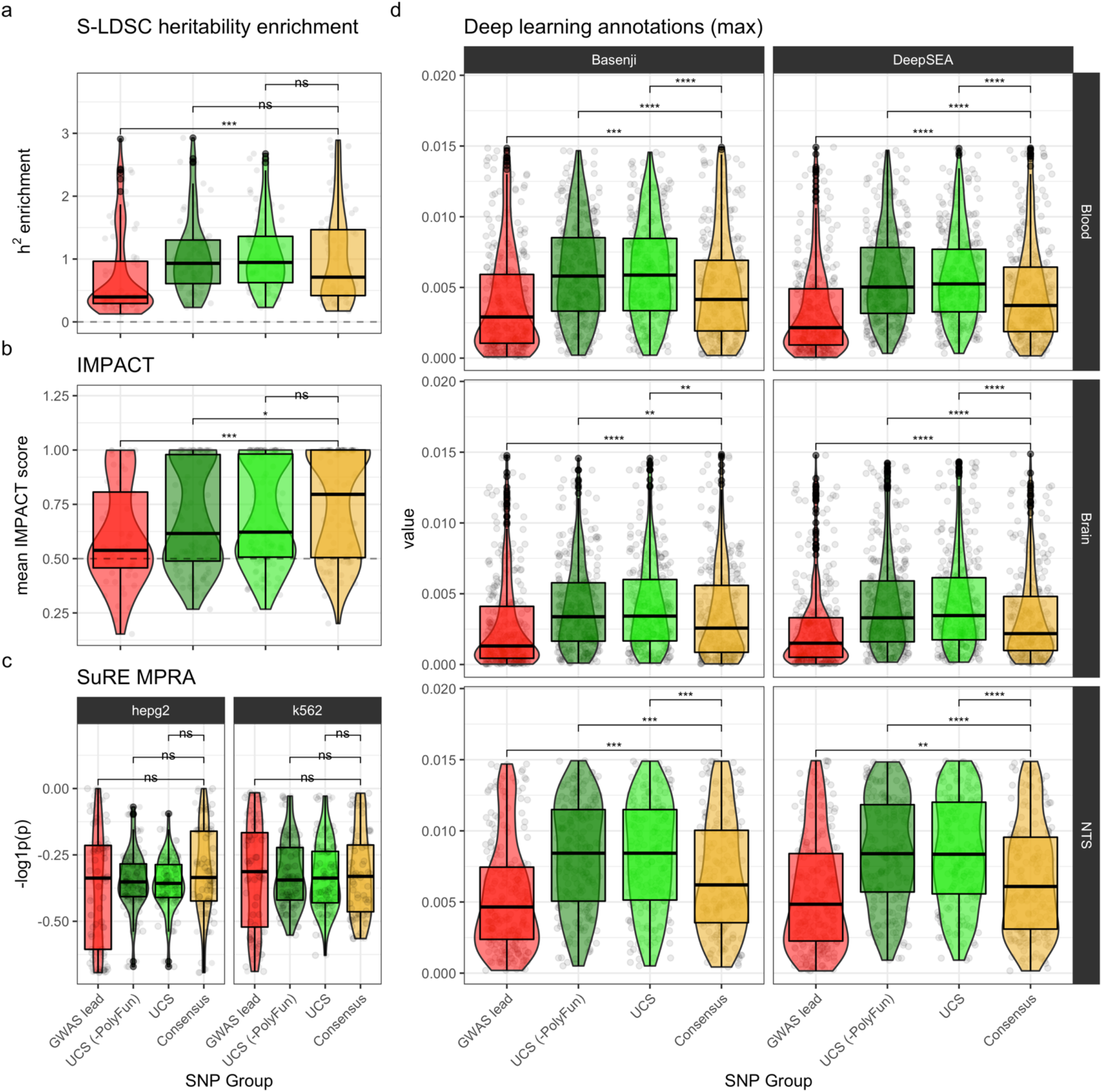
*In silico* validation. *In silico* validation was performed by statistically comparing each SNP group. For each validation (panel), per-locus mean values were computed for each SNP group (each represented by a point). Significance labels are from Holm-adjusted p-values of pairwise Wilcoxon rank-sum tests. *Significance key: ns: p > 0*.*05, *: p ≤ 0*.*05, **: p ≤ 0*.*01, ***: p ≤ 0*.*001, ****: p ≤ 0*.*0001*. **a**, Mean PolyFun-implemented^17^ S-LDSC^39–41^ heritability (h^2^) enrichment per locus per SNP group. **b**, For each locus, the annotation with the highest mean IMPACT score across all SNPs in that locus was selected (to reduce noise from less relevant annotations). **c**, SuRE MPRA results. Negative log_10_-transformed mean p-values from Wilcoxon rank-sum tests comparing gene expression changes between reference and alternative allele genotypes. Cell lines (i.e. HepG2 and K562) are separated into columns. **d**, Deep learning model predictions of the epigenomic impact that mutating each genomic position would have. Tissue-specific models (i.e. Blood, Brain) were trained on datasets from only that tissue. Non-tissue-specific (NTS) models were also trained using all tissues together. For the purpose of readability, we collapse values across assays and remove outliers above the 75% percentile (see **Fig. S4** for full view).

## Discussion

Using a suite of complementary fine-mapping tools and validating using multiple independent lines of evidence we identified 1-4 high-confidence consensus SNPs in most known PD-associated loci. These Consensus SNPs had a high probability of being causal in PD, often through the disruption of regulatory elements. Colocalization tests and brain cell-type-specific chromatin accessibility and histone modifications allowed us to explore the locus-specific mechanisms by which the Consensus SNPs exert their effects. We showed that PD risk alleles in Consensus SNPs likely alter the functioning of microglia-specific enhancers in the loci LRRK2 (**Fig. 3a**) and FCGR2A (**Fig. 4b,d**), specifically through disrupting a *SPIB* binding motif in the latter. We also identified Consensus SNPs affecting an oligodendrocyte-specific *MBNL2* enhancer (**Fig. 3b**) and a neuron-specific *DYRK1A* enhancer (**Fig. 3a,c**), the latter of which is mediated through altered binding with *PATZ1*. Additionally, in two loci (KCNIP3, TRIM40) our Consensus SNPs were missense mutations, both of which were found to contain missense mutations (2 and 6, respectively) and the former of which was significant in a Bonferroni-corrected rare coding variant burden analysis^4^.

The fine-mapped SNPs identified in this study consistently demonstrate higher functional relevance than GWAS lead SNPs according to multiple lines of evidence, including h^2^ enrichment, predictions from a suite of high-accuracy machine-learning models, cell-type-specific epigenomic assays and high-throughput variant-editing experimental assays. In particular, Consensus SNPs have the advantages of high functional enrichment, high coverage across all loci (100%), and relatively small set sizes (1-4 SNPs per locus), making them ideal candidates for further experimental validation. This remains true even after multiple testing correction, rigorous bootstrapped testing regimes, and removing any functionally informed fine-mapping results during UCS identification to avoid circularity in our validation strategy.

Together, these multiple lines of evidence support our hypotheses that high-quality fine-mapped SNPs are much more likely to be causal for PD than simply taking the SNPs with the smallest p-values, or random SNPs.

The present study has several limitations and could further be improved upon in the future by addressing the following: 1) our LD reference panels did not come from the same participants as the original PD GWAS studies, which reduces the accuracy of fine-mapping^20^, 2) not all fine-mapped variants are mediated via cell-type-specific enhancers, and thus future studies would benefit from exploring other mechanisms not explored here such as splicing (e.g. in MAPT)^47,48^, 3) the PD GWAS was conducted using almost entirely genotyping data (as opposed to whole-genome sequencing), which can introduce substantial bias and miss rare variants as a consequence of imputation procedures, 4) the functional genomic annotations used in this study are far from a complete representation of all PD-relevant tissues, brain regions (e.g. substantia nigra), cell-types, and assay modalities across varying PD stages and subtypes, and 5) functional experiments would further validate these results. Even so, we hope these results will continue to be a resource for future studies with better powered, more diverse datasets across a variety of tissues, cell-types, and physiological conditions.

In summary, we have fine-mapped almost all known PD-associated loci using a suite of complementary fine-mapping methods and identified putative mechanisms of actions through which they increase risk of PD, including cell-type and regulatory element type. Furthermore, our consensus fine-mapping strategy, implemented in *echolocatoR*, can easily be applied to any other GWAS/QTL summary statistics, opening up many opportunities for rapid and robust identification of causal genetic variants. This sets the stage to further our understanding of PD through uncovering potential shared genomic mechanisms underlying both PD and other neurological diseases, such as Alzheimer’s disease. Lastly, all SNP-wise fine-mapping results, LD matrices, merged annotations and plots from this study have been made available through a dedicated web application (https://rajlab.shinyapps.io/Fine_Mapping_Shiny), opening many opportunities to explore each locus in greater depth and, importantly, to validate putative causal SNPs through experimental validation (e.g CRISPR-cas9 editing in patient-derived iPSC models).

## Methods

All variant-level annotations, tools and analyses used in our pipeline were integrated into *echolocatoR^11^*, either directly or via APIs, and run in R v3.6.3. See **Supplementary Methods** for a more detailed description of each dataset.

### GWAS

Full genome-wide GWAS summary statistics from Nalls and colleagues^4^ were provided by the authors, and by 23andMe. If unspecified, we identified the lead GWAS SNP as the one with the lowest corrected p-value within that locus. Then, for each locus we gathered all SNPs within 2MB windows (i.e. ± 1Mb flanking the lead GWAS SNP) and filtered out SNPs with a minor allele frequency (MAF) < 0.05. We focused on common variants in order to maximize the relevance of these results to a larger proportion of the PD population.

### Fine-mapping

Statistical fine-mapping was performed on each locus separately with ABF^13^, FINEMAP^13–15^ and SuSiE^16^. Functional fine-mapping was performed using PolyFun+SuSiE^16,17^, which computes SNP-wise heritability-derived prior probabilities using a L2-regularized extension of stratified LD SCore (S-LDSC) regression^39–41^. For PolyFun+SuSiE, we used the default UK Biobank baseline model composed of 187 binarized epigenomic and genic annotations^49^. In all subsequent analyses presented here, loci that fall within the HLA region or the Tau/17q21.31 region were excluded due to the particularly complex LD architectures^12^.

While the specifics of each fine-mapping model differ from tool-to-tool, they are united by several key features: 1) they are Bayesian models, 2) they provide the PP that each SNP is a causal SNP, on a scale from 0 to 1, 3) they provide CS of SNPs that have been identified as having a high PP of being causal, which we have set at a threshold of PP ≥ 0.95 for all tools (see **Supplementary Methods** for details). We only included tools that met the following criteria: 1) can take into account LD, and 2) can operate using only summary statistics, which are more widely accessible and perform comparably to models using individual-level genotype data^5^. By running all of these tools on each PD locus, we reduced the average number of candidate SNPs per locus from ~5,000 to 8 (**Fig. 2**). For all fine-mapping models we set the (maximum) number of causal SNPs to five, except for ABF which can only assume a single causal SNP.

### LD

LD correlation matrices (in units of *r*) were acquired for each locus from the UK Biobank (UKB) reference panel, precalculated by Weissbrod et al.^17^. We also acquired LD from the 1000 Genome Project (1KG), both Phases 1 and 3^50^, by downloading the relevant variant call files (VCF) from the 1KG file transfer protocol (FTP) server using Tabix^51^ and then using the R package *snpStats^52^* set to the default parameters to calculate all pairwise *r* values. For both the UKB and 1KG reference panels, individuals of non-European ancestry were removed to best match the populations in the PD GWAS data. Any SNPs that could not be identified within the LD reference were necessarily removed from subsequent analyses.

### eQTL

*echolocatoR^11^* uses another R package developed by our lab, *catalogueR*, to automatically query all 110 eQTL datasets (from 20 studies) in the *eQTL Catalogue^53^* via an API. This comprised the majority of the eQTL datasets used in this study as they have all been uniformly re-analyzed using the exact same pipeline with standardized formatting. This includes data from the Genotype-Tissue Expression (GTEx) project V8, including 13 brain regions: Amygdala, Anterior cingulate cortex (BA24), Caudate, Cerebellar Hemisphere, Cerebellum, Cortex, Frontal Cortex (BA9), Hippocampus, Hypothalamus, Nucleus accumbens, Putamen, Spinal cord (cervical c-1), and Substantia nigra. We also obtained genotype gene expression data from the Multi-Ethnic Study of Atherosclerosis (MESA) ^54^, which contains eQTL data in monocytes from multiple subpopulations: 233 African American (AFA), 578 European (CAU) and 352 Hispanic (HIS) individuals.

### Colocalization

A GWAS-QTL locus pair was considered “colocalized” if the following criterion were met, where the posterior probability that the signals are associated with their respective traits and shared is PP.H4 and the posterior probability that the signals are associated with their respective traits but *not* shared is PP.H3):

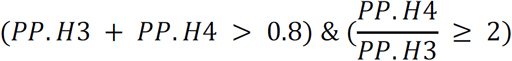

This provides a more robust means of identifying whether the signal in a GWAS dataset can be considered the same as the signal within a QTL dataset, as opposed to simply looking at the number of overlapping significant QTLs which can be confounded by factors such as LD^55^.

### Transcription factor binding motifs

We ran *motifbreakR^35^* on select Consensus SNPs, which 1) identifies whether a given variant falls within binding motifs or one or more transcription factors (TFBM) and 2) calculates how much each allele of that variant impacts motif binding. Motifs are compiled by a supporting package, *MotifDb*, which contains a comprehensive database of positional weight matrices (PWM) (n = 9,933) from 14 TFBM databases gathered from multiple organisms (n = 16 species) and assay types (including ChIP-seq). For each variant, *motifbreakR* first queries the PWM databases to identify any motifs that the variant may fall within, and returns metrics assessing how much each allele of that variant disrupts (or enhances) binding in the putative TFBM(s). Particularly important metrics include *pct_ref/_alt*: the proportion of maximum binding affinity of the motif to a given TF (0 to 1 scale) afforded by the variant’s reference/alternative alleles (respectively), and *allele diff*.: The difference in the proportion of binding affinity between the reference (pct_ref) and alternative (pct_alt) alleles (-1 to 1 scale). Next, *motifbreakR* applies a robust background correction regime to compute p-values. This is a computationally expensive but nevertheless important step as it helps guard against false positives, a substantial concern as many motifs have similar sequences and/or promiscuously bind to multiple transcription factors. Lastly, we applied the Benjamini–Hochberg procedure to compute false discovery rate (FDR) to account for querying multiple variants^56^.

### *In silico* validation

S-LDSC heritability, IMPACT, SuRE MPRA, and deep learning annotations

For all S-LDSC heritability scores, IMPACT scores, SuRE MPRA p-values, and deep learning epigenomic predictions (collectively referred to as validation annotations), the we calculated the mean of the respective values per locus per SNP group (*GWAS lead, UCS (-PolyFun), UCS, Consensus*). Mean was used as opposed to max values, to avoid bias due to differences in SNP group sizes. We evaluated independence between SNP groups using a series of pairwise Wilcoxon rank-sum tests with multiple-testing adjusted p-values (Holm-Bonferroni method) (**Fig. 5**).

To ensure the robustness of these results, we also employed a bootstrapping hypothesis testing procedure, analogous to that proposed by Tibshirani and Efron^57^. For 10,000 iterations in each SNP group, repeated separately for each validation annotation, we sampled (with replacement) 20 per-locus validation annotation means of the SNP group of interest and used a Wilcoxon rank sum test (*coin::wilcox_test* in R) to determine whether their validation annotation values significantly differed from that of 20 randomly sampled SNPs (*Random*). Importantly, our *Random* samples were taken only from the 2Mb windows defining our PD loci (as opposed to the whole genome), as an appropriate background to which compare the SNP groups. Resampling 10,000 times ensures that these results are not merely due to chance. Finally, we used a generalized linear model (using the R *stats::glm* function) to test whether the normalized test statistic distribution (*z-value*) of each fine-mapped SNP group (*UCS (-PolyFun), UCS, Consensus (-PolyFun), Consensus*) was significantly different from the *GWAS lead* z-value distribution. This bootstrapping procedure can easily be replicated using a single *echolocatoR* function, *VALIDATION*.*bootstrap*.

Lastly, we repeated enrichment tests for each SNP group against each combination of bulk brain tissue and cell-type-specific epigenomic peaks, regulatory elements, and interactomes (**Fig. S6**).

### Cell-type-specific epigenomic annotations

We conducted a series of negative binomial enrichment tests, with the *XGR::xGRviaGenomicAnno* function in R^58^, using the following annotations: 1) cell-type-specific epigenomic peaks (scATAC, ATAC, H3K27ac, H3K4me3), 2) cell-type-specific regulatory regions (enhancers, promoters), 3) cell-type-specific interactome anchors (PLAC, cicero), 4) bulk brain epigenomic peaks (ATAC), 5) bulk brain interactome anchors (HiChIP_FitHiChIP)^18,19^. We conducted these tests separately for each combination of assay/regulatory element and cell-type. Furthermore, enrichment tests were repeated separately for each SNP group. All SNPs within any 2Mb locus were used as the background.

These same epigenomic datasets were used in summary plots (**Fig. 2**) and track plots (**Fig. 3-4**). Annotations for missense variants were gathered from *biomaRt* ^59,60^ and *HaploReg* ^61^.

## Data Availability

A significant subset of the PD GWAS summary statistics can be found in the supplementary materials of the original publication^4^. For full summary statistics, please contact the respective authors of that publication. *eQTL Catalogue* data is freely accessible through the main website (https://www.ebi.ac.uk/eqtl) or through the *catalogueR* software (https://github.com/RajLabMSSM/catalogueR).

All results of this study (fine-mapping, colocalization, plots) are accessible through the *Fine-mapping Results Portal*, which the authors have created to easily share and visualize the results of this study and others by our lab (https://rajlab.shinyapps.io/Fine_Mapping_Shiny).

## Code Availability

R scripts containing all of the analyses conducted in this study are also made freely available on GitHub: https://github.com/RajLabMSSM/Fine_Mapping

Code for the *Fine-mapping Results Portal* is also available on GitHub: https://github.com/RajLabMSSM/Fine_Mapping_Shiny

*echolocatoR* and *catalogueR* are an open-source R packages that can be installed through the following GitHub repositories:

https://github.com/RajLabMSSM/echolocatoR

https://github.com/RajLabMSSM/catalogueR

## Acknowledgments

We would like to thank Kuan-lin Huang, Jack Humphrey, Ricardo Vialle, Elisa Navarro, Giulietta Riboldi, Gloriia Novikova, Cecilia Lindgren and Teresa Ferreira for their valuable feedback and suggestions. We would also like to thank Omer Weissbrod, Christopher Glass, Alexi Nott for their guidance with data and/or tool integration. This work was supported by grants from Michael J. Fox Foundation (MJFF) and the US National Institutes of Health (NIH NIA R01-AG054005 and NIA R56-AG055824). This work was supported in part through the computational and data resources and staff expertise provided by Scientific Computing at the Icahn School of Medicine at Mount Sinai. We thank the research participants and employees of 23andMe who contributed to the PD GWAS.

## A note from the first author (BMS)

This study is dedicated to Robert Neil Cronin, my grandfather. He is the reason I decided to pursue this field as a career, has continued to be my source of motivation, especially when challenges arose, and has kept this work grounded in its core mission of helping others. Without him, this study as it exists today would not have been possible. For all of this, and so much more, I am eternally grateful.

## Supplementary Materials

### URLs

Fine-mapping Results Shiny App: https://rajlab.shinyapps.io/Fine_Mapping_Shiny

Fine-mapping Results GitHub: https://github.com/RajLabMSSM/Fine_Mapping

echolocatoR: https://github.com/RajLabMSSM/echolocatoR

catalogueR: https://github.com/RajLabMSSM/catalogueR

PolyFun: https://github.com/omerwe/polyfun

FINEMAP: http://www.christianbenner.com

S-LDSC: https://github.com/bulik/ldsc

SuSIE: https://stephenslab.github.io/susieR/index.html

coloc: https://chr1swallace.github.io/coloc/index.html

eQTL Catalogue: https://www.ebi.ac.uk/eqtl

The 1000 Genomes Project FTP server: http://ftp.1000genomes.ebi.ac.uk

UKB LD matrices (pre-computed by the Price Lab): https://data.broadinstitute.org/alkesgroup/UKBB_LD

Single Cell Portal: https://singlecell.broadinstitute.org/single_cell

IMPACT: https://github.com/immunogenomics/IMPACT

Basenji: https://github.com/calico/basenji

DeepSEA: https://hb.flatironinstitute.org/deepsea

### Supplementary Methods

Averaged (across loci) mean (across fine-mapping tools) PP was calculated per SNP group : lead GWAS SNPs (1 SNP/locus, PP = 0.533), UCS (~8 SNPs per locus, PP = 0.351), Consensus SNPs (~3 SNPs per locus, PP = 0.575).

#### Fine-mapping

ABF was tested across all loci through the use of the *coloc*.*abf* function within the *coloc* R package^55^. ABF can only assume a single causal SNP, and thus can’t model scenarios where this is more than one true causal SNP, or for that matter, scenarios where two or more SNPs are indistinguishable due to perfect LD. SNPs with PP ≥ 0.95 were considered part of the ABF credible set, and added to the overall cross-tool Credible Set. However we decided to exclude this method when identifying Consensus SNPs because it almost exclusively returned the lead SNP, suggesting this simpler algorithm was not offering any additional information beyond which SNP had the lowest GWAS p-value. SuSiE was implemented via the *susieR* R package^16^ by using *susie_bhat* according to the parameters recommended in the documentation for fine-mapping based on summary statistics, except for *L* (max causal variants) which was set to five. For each locus, SuSiE could produce multiple credible sets, which within the context of this tool particular tool are defined as sets of SNPs that has a ≥ 95% PP of containing one or more causal SNPs, but was unable to distinguish which of these SNPs are truly causal due to very high or perfect LD (*r*^2^ ≈1). All SNPs from each SuSiE credible set were added to the *echolocatoR* Credible Set, and their PP were compiled.

PolyFun+SuSiE was run using the pre-computed heritability-derived priors from 15 UKB GWAS traits that come with the PolyFun software ^17^, as this multi-trait heritability was predicted to be translatable to other traits. We also computed heritability using PolyFun’s modified version of S-LDSC^39–41^ with the PD GWAS and found negligible differences in the resulting posterior probabilities.

We set the parameters for FINEMAP as recommended by the documentation: *--model cond* (to fine-map using stepwise conditional search, as opposed to stochastic shotgun search), *--corr-config 0*.*95, --corr-group 0*.*99, --n-configs-top 50000, --prior-k 0, --prior-k0 0, --prior-std 0*.*05*.FINEMAP produces two main output files: 1) *data*.*config*, which contains different configurations of credible sets and their PP of including the causal SNP(s). Only the configuration with the highest PP was used, and 2) *data*.*snp*, which contains SNP-wise PPs that each SNP is causal.

#### Annotations

The *biomaRt* R package (Durinck et al., 2005; Durinck, Spellman, Birney, & Huber, 2009) was used to download SNP-level annotations for the following fields: *ensembl_gene_stable_id, consequence_type_tv, polyphen_prediction, polyphen_score, sift_prediction, sift_score, reg_consequence_types*, and *validated*.

The HaploR R package (Zhbannikov, Yashin, Arbeev, & Ukraintseva, 2017) was used to query the HaploReg and RegulomeDB databases for the following SNP-level annotations: Reference allele frequencies within the African American (*AFR*), American (*AMR*), Asian (*ASN*), and European (*EUR*) subpopulations, *GERP_cons, SiPhy_cons, Chromatin_State, Chromatin_States_Imputed, Chromatin_Marks, DNAse, Proteins, eQTL, gwas, grasp, Motifs, GENCODE_id, GENCODE_name, GENCODE_direction, GENCODE_distance, RefSeq_id, RefSeq_name, RefSeq_direction, RefSeq_distance, dbSNP_functional_annotation, query_snp_rsid, Promoter_histone_marks, and Enhancer_histone_marks*.

#### Colocalization of GWAS and QTL signals

We first compiled expression quantitative trait locus (eQTL) datasets from the *eQTL Catalogue^53^*, 13 brain tissues from GTEx V7^64^, and Multi-Ethnic Study of Atherosclerosis (MESA)^54^. In total, we included 110 eQTL datasets from 21 different studies. Colocalization tests^55^ were conducted between each PD GWAS and each overlapping eQTL locus (repeated for each eGene) to determine whether these trait-associated signals were shared across datasets. Of the resulting 155,577 tests, 546 were colocalized according to standard criterion (see Methods). We focused on 188 comparisons that had extremely high colocalization probability (> 0.95) to reduce the risk of false positives (**Fig. S1**). Of these high-probability GWAS-QTL loci, genome-wide significant eVariants overlapped with lead GWAS SNPs in 10 loci, UCS SNPs in 47 loci, and Consensus SNPs in 31 loci. This pattern of results indicates that, relative to simply using the lead GWAS SNPs, *echolocatoR* identifies SNPs with a substantially higher probability of having causal impacts on gene expression, specifically within relevant tissues and cell types.

## Supplementary Figures

Fig. S1. | Comparisons of fine-mapping results between LD panels and studies.

**a-f**, Pairwise Spearman’s rank-order correlations between fine-mapping results using either UKB or 1KG LD reference panels. As expected, there is a perfect correlation when using ABF as this tool does not incorporate LD information. The SNP-wise posterior probabilities from SuSiE and PolyFun+SuSiE are very consistent between LD panels (Spearman’s rho = 0.73 and 0.76, respectively). In contrast, FINEMAP shows no significant relationship between the results with each LD panel, calling into question its robustness to LD misspecification. Multi-tool mean PP showed slightly weaker, but still highly significant, correlation between LD panels (Spearman’s rho = 0.65).

**g**, Heatmap of all pairwise Spearman’s rank-order correlations between each fine-mapping tool and LD panel combination.

**h-m**, Pairwise Spearman’s rank-order correlations between SNP-wise posterior probabilities generated by each of the fine-mapping methods used in this study with those of Grenn et al. (generated using FINEMAP)^21^. Our FINEMAP results, as well results from the other tools we used (ABF, SuSiE, PolyFun+SuSiE) to a lesser degree, showed a strong and significant correlation with those of Grenn et al (Spearman’s rho = 0.68, p < 2.2×10^-16^).

Fig. S2. | All colocalized PD GWAS-eQTL loci.

Only loci that had a > 95% probability of colocalization (using *coloc*) are shown here. Shape markers indicate that there was overlap between a genome-wide significant eVariant within a given eQTL locus (p < 6.38×10^-11^) and ≥ 1 SNP from the following SNP groups: lead GWAS SNP (black square border), Union Credible Set SNP (cyan diamond), Consensus SNP (cyan dot). eQTL datasets are grouped by tissue system: Blood, central nervous system (CNS) and Other.

Fig. S3. | Extended *motifbreakR* results and single-cell gene expression.

The following results were obtained through the Single Cell Portal:

**d**, *PATZ1* is expressed in neurons but not glia. Data comes from mouse hippocampus^65^.

**e**, *DYRK1A* is expressed in multiple brain cell-types, but primarily neurons. Data comes from mouse hippocampus^65^.

**f**, *FCGR2A* is expressed in macrophages, including microglia. Data comes from human melanoma samples^66^.

**g**, *SPIB* is most strongly expressed in glia, and especially microglia. Data comes from mouse visual cortex^67^.

Fig. S4. | *In silico* validation: extended SNP groups.

In silico validation in an extended list of SNP groups: 1) *Random*: randomly selected SNPs (three per locus to approximate the Consensus SNP set size), 2) *GWAS sig. nom*.: nominally significant PD GWAS SNPs (p < 0.05), 3) *GWAS sig*.: genome-wide significant PD GWAS SNPs (p < 5×10^-8^), 4) *GWAS lead*: the SNP with the smallest p-value (if multiple SNPs had the same lowest p-value, the one with the largest effect size was taken), 5-8) *<tool> CS*: fine-mapping tool-specific CS_95%_ SNPs, 9) *UCS*: Union Credible Set SNPs, 10) *Consensus*: SNPs in the CS_95%_ of at least two fine-mapping tools, and 11) *Consensus (-POLYFUN_SUSIE)*: Consensus SNPs called after excluding PolyFun+SuSiE results to guard against circularity (as ENCODE and Roadmap data was used to train both PolyFun and many of the models we use here for validation).

Fig. S5. | *In silico* validation: bootstrap analysis plots.

For each bootstrapping analysis, we plotted the normalized test-statistic distributions (z-statistic) from 10,000 iterations of Wilcoxon rank-sum tests (per SNP group) and applied kernel density estimation for smoothing. The labels at the peak of each distribution display the p-value from a generalized linear model (GLM) testing for independence between the *GWAS lead* z-statistic distribution and that of each fine-mapped SNP group (e.g. UCS, Consensus).

Fig. S6. | Cell-type-specific epigenomic enrichment.

Results from a series of enrichment tests (*XGR*) checking for overlap between each SNP group and cell-type-specific epigenomic assay peaks (H3K27ac, H3K4me3, and scATAC), cell-type-specific regulatory regions (enhancers and promoters), cell-type-specific scATAC-seq co-accessibility anchors (cicero), as well as epigenomic peaks from bulk brain tissue (ATAC). *nOverlap* indicates the number of SNPs overlapping with the respective annotation. Log_10_(x+1) fold-change (fc; *x-axis*) is plotted against the multiple-testing corrected false discovery rate (FDR; *y-axis*). Best fit lines from grouped linear regression models (y ∼ x for each SNP group) are shown with confidence intervals. The horizontal dashed line indicates the FDR ≤0.05 threshold.

## Supplementary Tables

Table S1.Consensus SNP summary across all loci.

1) Novel loci identified in Nalls et al. (2019) are listed at the top,

2) Highest to lowest posterior probability (PP) summed across fine-mapping methods and averaged across Consensus SNPs (PP.mean.sum),

3) highest to lowest number of QTLs, averaged across Consensus SNPs (QTL.mean.count),

4) lowest to highest number of Consensus SNPs (SNP.count).

Also included is whether each Consensus SNP is the lead SNP from the Nalls et al. (2019) PD GWAS (Y = Yes, N = No).

