## Supplementary figures and images for "Fine-mapping of Parkinson’s disease susceptibility loci identifies putative causal variants"

### Fig. S1

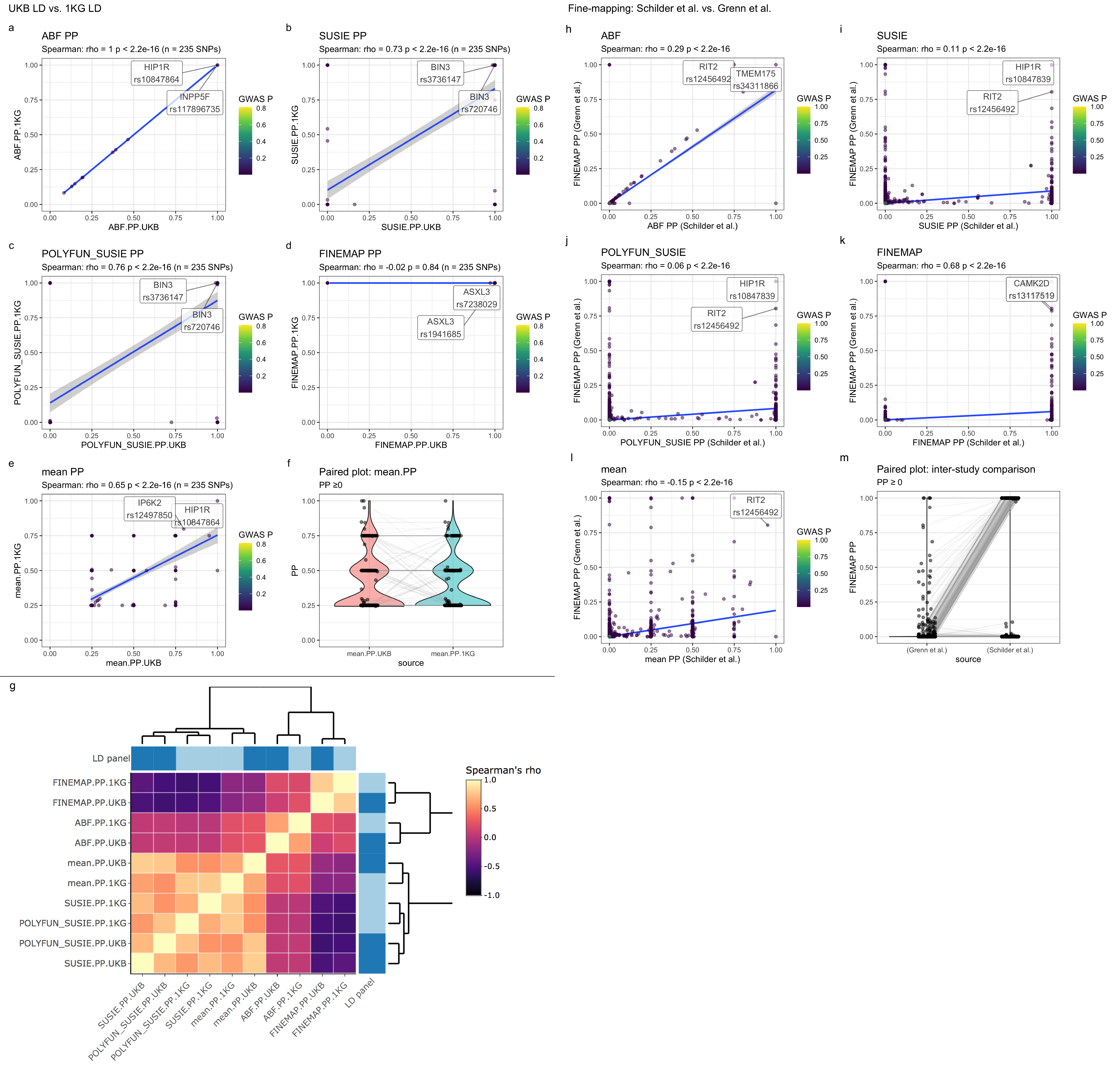

### Fig. S2

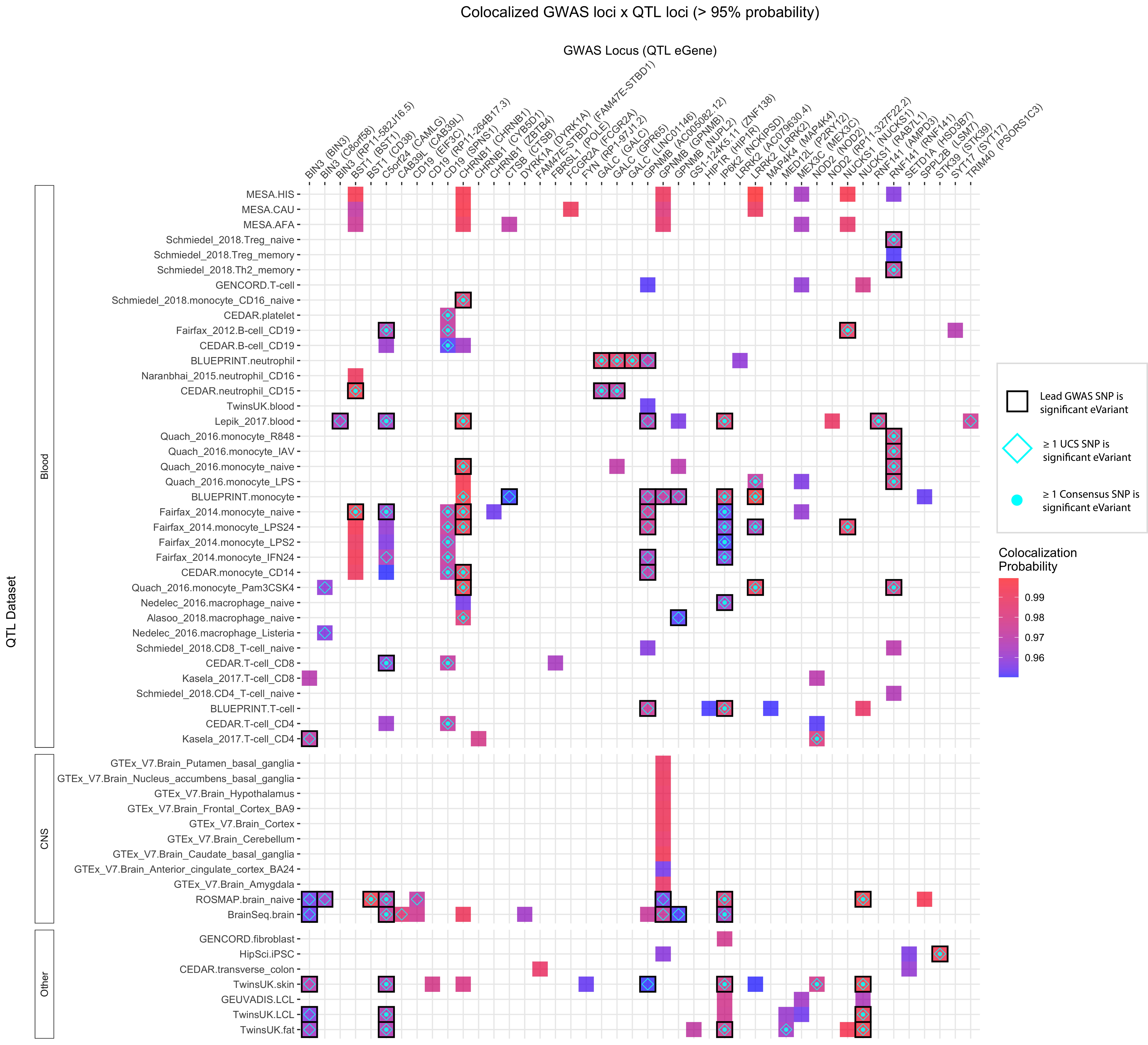

### Fig. S3

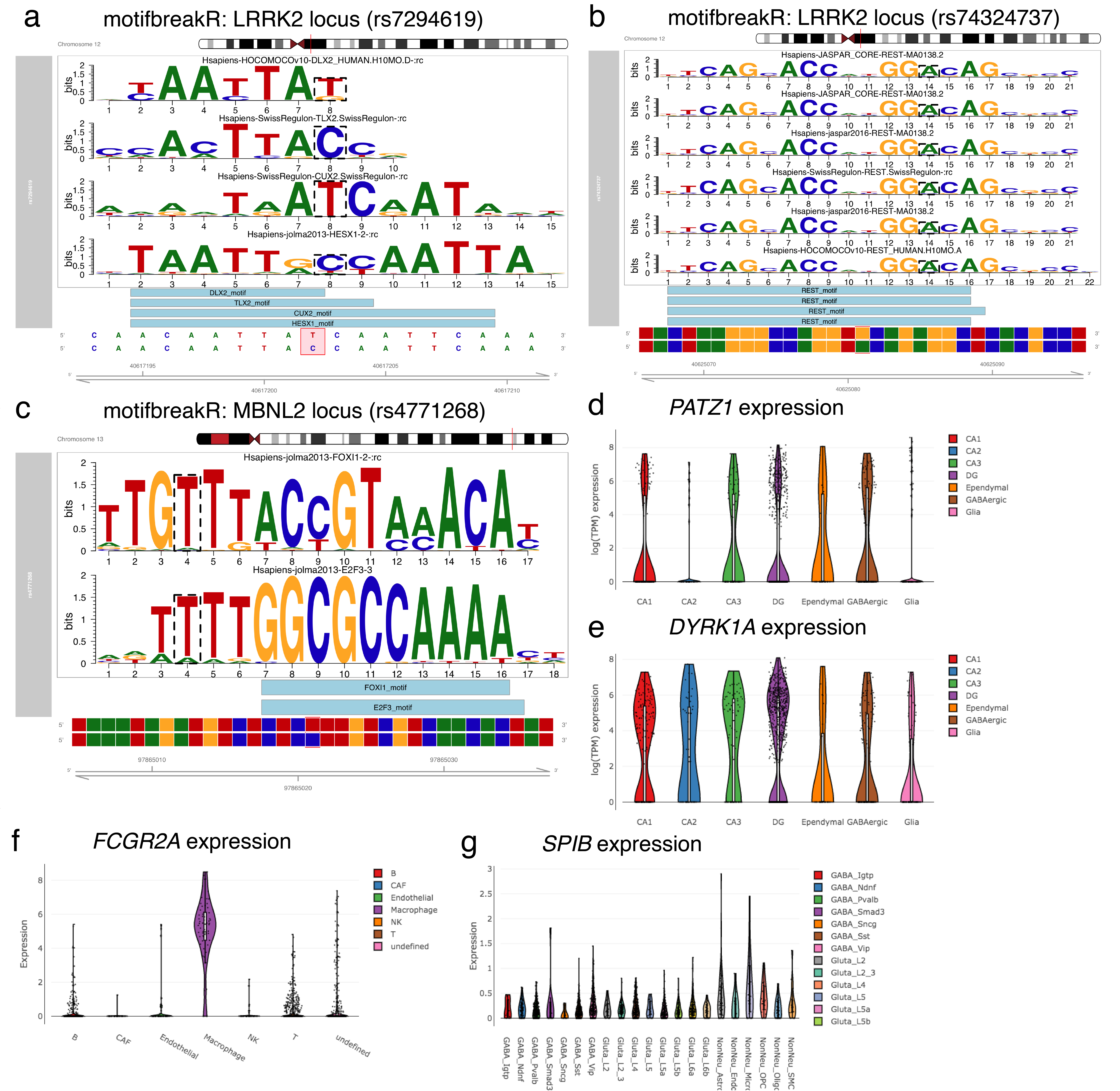

### Fig. S4

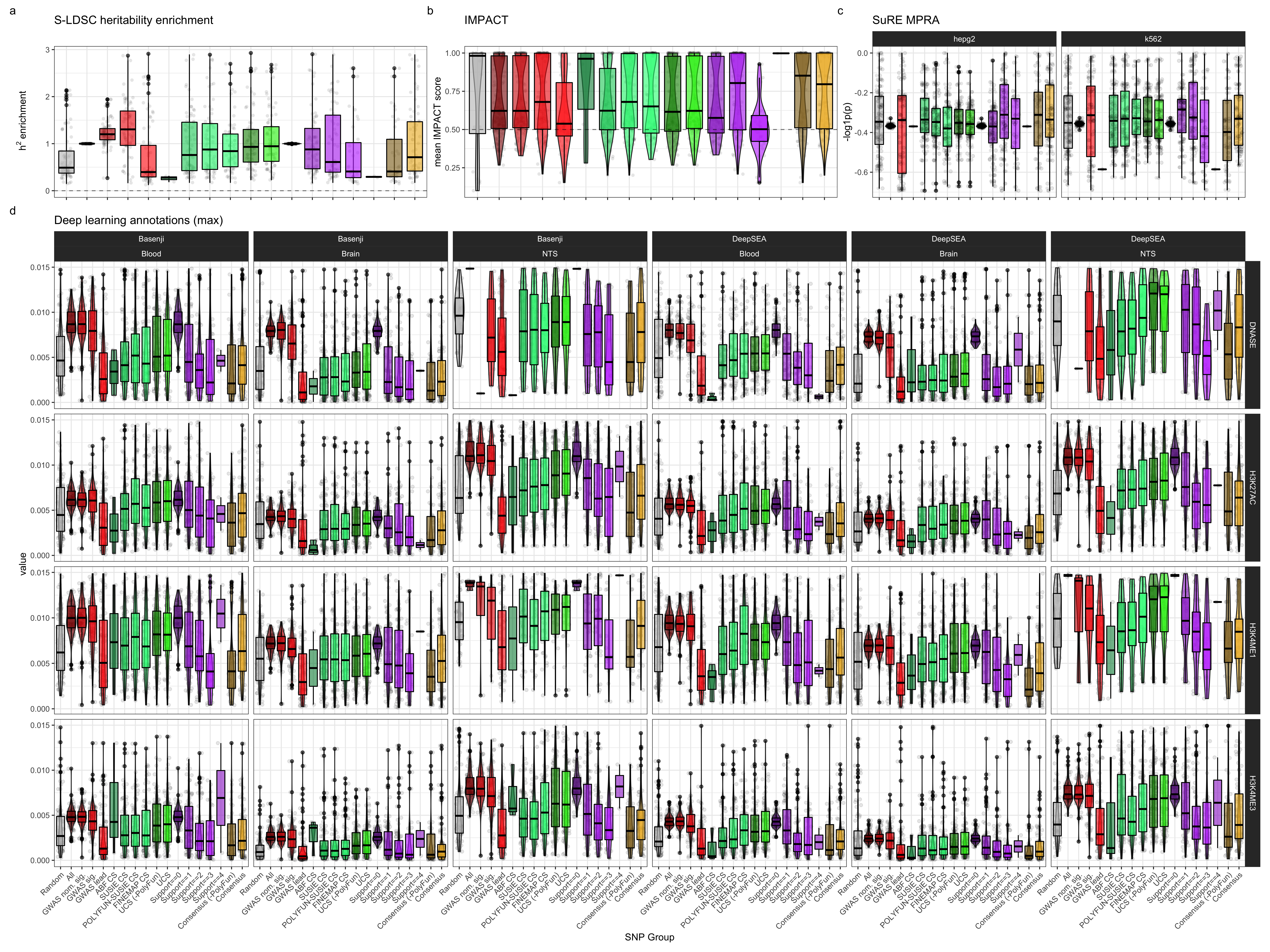

### Fig. S5

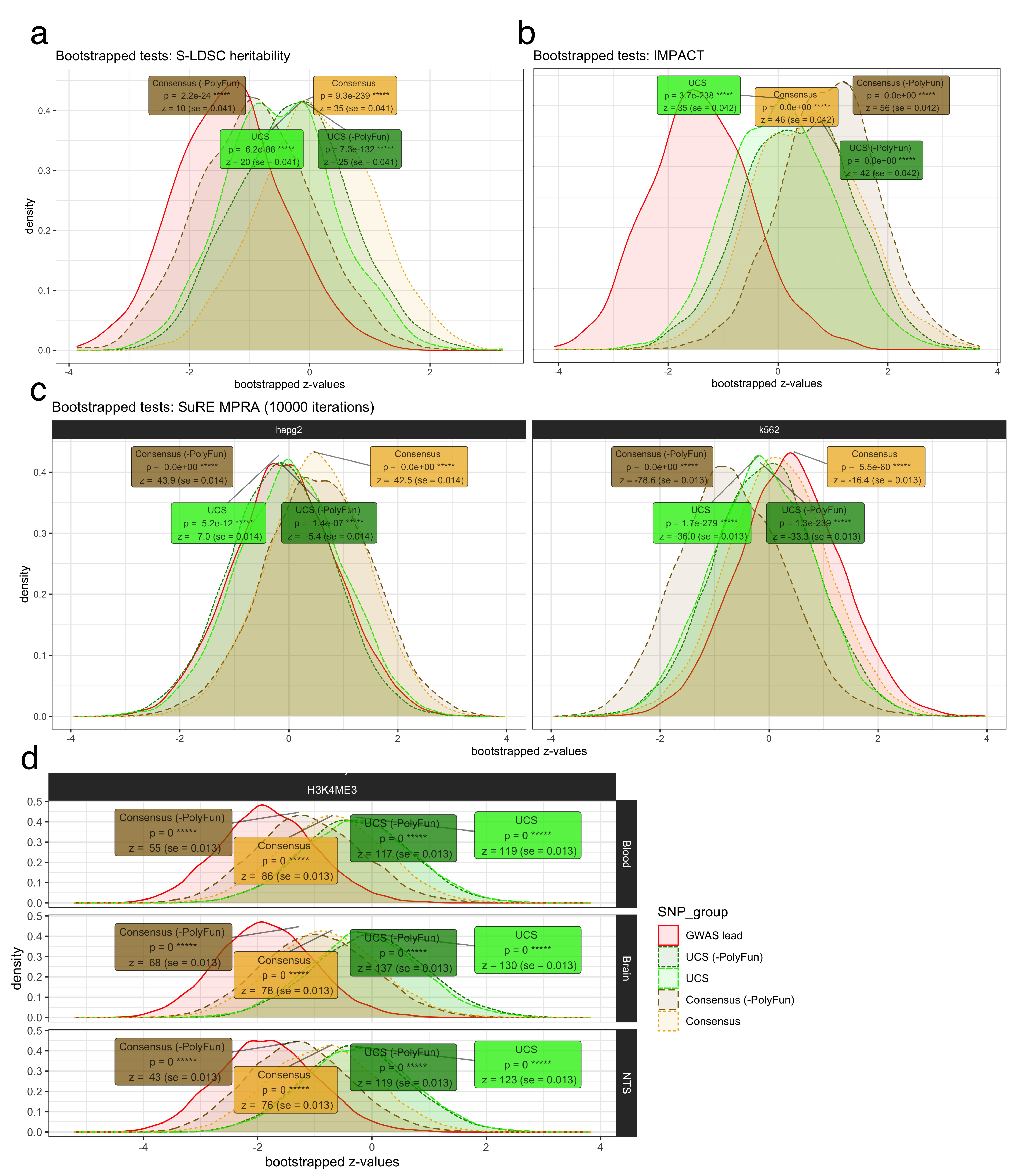

### Fig. S6

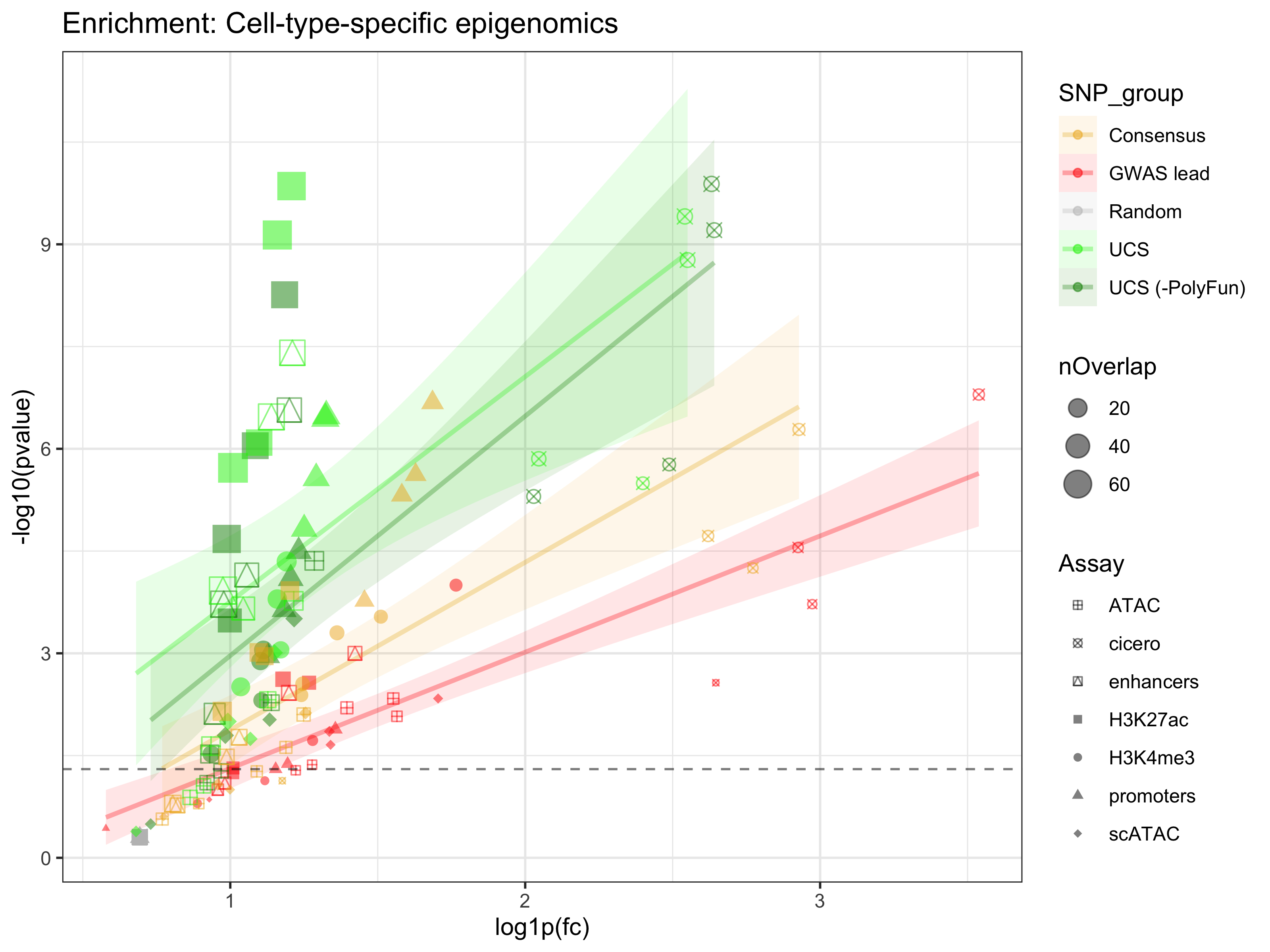
